# Increased surface area of Heschl’s gyrus following music-cued motor training

**DOI:** 10.1101/2025.10.09.675791

**Authors:** Mohammed A. Mudarris, Emma Moore, Neil Roberts, Mark E. Bastin, Dieuwke S. Veldhuijzen, Serge A. R. B. Rombouts, Katie Overy, Rebecca S. Schaefer

**Author notes:** Corresponding Author* M. A. Mudarris.

## Abstract

Previous work found that adding musical cues to a four-week, left-handed, visuomotor sequence training task increased white matter connectivity between contralateral auditory and motor regions. In this study, grey matter measures are investigated by examining changes in cortical auditory and motor regions, specifically the superior and transverse temporal gyri, precentral gyrus, and paracentral lobule. Thirty healthy young adults trained on left-handed finger sequences over four weeks, either with visual cues only (Control group) or with added musical cues (Music group). Participants underwent MRI and behavioral assessments pre- and post-training. Results revealed evidence of a time-group-hemisphere interaction, showing increased surface area of the contralateral, right Heschl’s gyrus in the Music group only, after training. Additional exploratory analyses revealed that behavioral performance was associated with surface area decreases in cortical motor areas in the Music group only, but these results did not survive multiple comparison correction. The results demonstrate auditory-specific neuroplastic changes related to adding musical cues to short-term visuomotor training in a controlled setting, furthering our understanding of patterns of neuroplasticity related to music-based movement, with implications for cued movement paradigms in health settings.

## Introduction

Over the past two decades it has been shown that acquiring a specific motor skill is associated with distinct structural neuroplastic changes. Additional to findings related to life-long training (e.g. Gaser & Schlaug, 2003), distinct neural plasticity has been shown longitudinally in novices following only weeks or months of motor training (e.g., Sale et al., 2017), and there is converging evidence that these changes involve both grey matter structures (Draganski et al., 2004; Lövdén et al., 2013; Sale et al., 2017), and white matter tracts (Blecher et al., 2016; Vaquero et al., 2018). When considering music training, such differences are not only seen for life-long professional musicians (e.g., Eierud et al., 2023; Gaser & Schlaug, 2003), but also for non-musicians who train on music-based movement (Bruchhage et al., 2020; Worschech et al., 2022). Besides instrumental music training, neuroplastic changes have also been reported for learning a new movement using music as a cue (Blecher et al., 2016; Moore et al., 2017; Vaquero et al., 2018). Cued movement is often used clinically, aiming to provide a temporal guide for movement, or to encourage more movement vigor through motivational mechanisms (e.g., Schaefer, 2014; Thaut, 2013). In a previous study, adding musical cues to a left-handed visuomotor training task was found to lead to significantly increased Fractional Anisotropy (FA) in the contralateral, right arcuate fasciculus (Moore et al., 2017), which is known to connect auditory and motor regions of the brain. Here, the aim is to explore whether this music-cued motor training leads to grey matter changes in specific auditory and motor regions.

Structural grey matter changes in auditory and motor regions have been reported in previous music-based motor learning paradigms. Specifically for music training, for instance, eight weeks of drum training led to cortical thickness increases in the paracentral lobule, connecting to both premotor and sensory areas, and superior frontal regions involved in processing of rhythmic temporal structure, and attending to a motor sequence (Bruchhage et al., 2020). When comparing six months of piano lessons with music listening as a control in a cohort of non-musician older adults, the piano training group showed significant increases in cortical thickness of the left Heschl’s gyrus, whereas the control group showed decreases, in both hemispheres, albeit stronger in the right hemisphere (Worschech et al., 2022). Additionally, in a study that included four weeks of finger-to-thumb motor sequence learning, but did not examine the role of music, decreased functional activation was observed in premotor and supplementary motor cortices for trained compared to novel sequences, along with increases in cortical thickness in primary and secondary motor areas and premotor regions, including pre- and post-central gyri (Sale et al., 2017). These findings underline the potential for plasticity of both motor and auditory grey matter areas for musical training over months, as well as for non-musically cued motor sequence training similar to the movement used here, over several weeks. Notably, these finding all report increases in cortical thickness, while very often, decreases for cortical areas are seen after learning a specialized task (Olivo et al., 2022; Wiestler & Diedrichsen, 2013). Specifically for learning a novel movement with musical cueing, to our knowledge, no results on grey matter changes have been previously reported.

These findings of change in auditory and motor area after musical training are also supported by findings on white matter connectivity, and specifically the relevance of the arcuate fasciculus, connecting auditory and motor regions, such as the finding that arcuate fasciculus volume and FA predict learning rate and learning speed in rhythm and melody tasks (Vaquero et al., 2018).

Moreover, how closely people move to the beat of a rhythm has been associated with the FA of left arcuate fasciculus (Blecher et al., 2016), and prior musical training was found to be associated with arcuate fasciculus volume asymmetry, showing greater volume in the right than the left hemisphere (Cui et al., 2024). For music-cued motor learning, increased FA in the arcuate fasciculus was found following a four-week motor sequence training with music and not without, and only in the hemisphere contralateral to the trained hand (Moore et al., 2017), indicating that these results of musical instrument training may also extend to music-cued motor learning.

Converging studies of the neuroanatomy of the arcuate fasciculus, using different modalities, confirm that it terminates posteriorly in the superior temporal gyrus, constituting the primary auditory cortex (A1; Maffei et al., 2015), and anteriorly at the inferior precentral gyrus, which encompasses primary and pre-motor areas (Brown et al., 2014). Given the demonstrated relevance of these areas for auditory-motor embedding in music-cued movement (Vaquero et al., 2021), the regions of interest in this study are thus focused specifically on two auditory regions, namely the Superior Temporal Gyrus (STG) and the Transverse Temporal Gyrus (TTG) together covering A1. We also examine two motor regions, namely the Precentral Gyrus (PCG) extending into the Paracentral Lobule (PCL) encompassing primary as well as supplementary motor regions, and reported to be closely related to PCG in the context of plasticity based on finger movements (Kimura et al., 2024).

Different MRI modalities tend to converge, with for instance cortical thickness correlating positively with white matter connectivity across the cerebrum (Gong et al., 2012). Arguably, grey matter structure, white matter connectivity, as well as functional activation during visual, auditory and motor tasks stem from the same underlying neuronal architecture (Hao et al., 2011). Therefore, it is also informative to investigate the degree of grey matter change related to white matter change. Different measures of grey matter are reported to be differentially related to white matter measures, where a change in one structural measure does not always correlate with another. Generally, larger grey matter surface area is thought to facilitate increased white matter connectivity, which in some cases is related to decrease in cortical thickness (Storsve et al., 2014), potentially through specialization (Olivo et al., 2022; Tremblay et al., 2021). Additionally, grey matter volume and cortical thickness seem to be closely associated and change together, differentiating from surface area (Storsve et al., 2014). These differential associations between white matter and specific measures of grey matter serve as the basis of the current investigation. Moreover, different aspects of grey matter have been shown to change with behavioral aspects of motor learning. However, here too, the changes in grey matter indices show mixed results. For instance, while some studies show an increase in cortical thickness with musical training (Bruchhage et al., 2020; Worschech et al., 2022), another study showed grey matter volume attenuates with a short finger tapping training (Olivo et al., 2022).

The aim of the current study is examine grey matter measures related to auditory and motor cortical regions, which have been previously linked to white matter connectivity, specifically increased FA of the arcuate fasciculus, following music-cued motor learning (Moore et al., 2017). Adding musical cues to motor training is expected to lead to changes in grey matter measures as compared to learning without musical cues, specifically in the auditory cortex, encompassed by the STG and TTG, as well as in primary-, and premotor regions, namely the PCG and PCL, all in the right hemisphere, contralateral to the trained hand, where the arcuate fasciculus was shown to be affected. Furthermore, this study explores how change in behavioral measures of sequence accuracy pre-and-post training may be related to changes in each of these cortical measures, considering that both increases and decreases related to learning have been reported. Finally, the study explores non-directional relationships between abovementioned cortical grey matter structures and our previously reported DTI findings on AF connectivity. The current paradigm is the first systematically controlled longitudinal investigation into cortical grey matter changes related to adding musical cues to finger movement training. Our findings aim to provide further evidence about music-related, learning-based plasticity in healthy young adults, and to increase our understanding of the neural mechanisms underlying motor learning. This will serve as a basis for further research examining the potential for plasticity and health-based applications using musical cueing in aging and clinical populations.

## Materials and Methods

### Participants

Thirty healthy young adults (18-30 years) participated in the study. Inclusion criteria consisted of right-handedness, as assessed with the Edinburgh Handedness Inventory (Oldfield, 1971), no history of neurological or psychiatric disorders, having had no more than six years of musical training, and not currently practicing a musical instrument. Participants were pseudo-randomized into either Music (*n* = 15; *M*age = 21.27, *SD* = 1.98 years; 4 males) or Control (*n* = 15; *M*age = 21.33, *SD* = 2.38 years; 5 males) groups. All participants gave informed written consent to participate and were monetarily compensated. The study was carried out in accordance with the Declaration of Helsinki and was approved by the local ethics committee of the University of Edinburgh and the West of Scotland Research Ethics Committee, UK (REC reference number 12/WS/0229).

### Motor Training

Participants trained on four different sequences, each comprising eight finger-to-thumb opposition movements, using the left (non-dominant) hand. Participants were asked to practice at home three times per week, for 20 minutes each, over four weeks. Both groups trained with visual cues presented in videos, showing circles descending along one of four vertical lines, with a horizontal line at the bottom of the screen signaling a pinching movement for the finger corresponding to that vertical line and the thumb. To ensure equal representation of the fingers across the four sequences, each sequence started with a different finger, and all four fingers were presented equally across the four sequences. Each sequence was comprised of eight movements indicated by the circle position. The Music group additionally had a soundtrack including a melody that contained temporal and pitch cues corresponding to each finger movement. The melody had a simple, steady beat consisting of eight notes (two bars), with a different melody per sequence, and a pitch matched to each finger.

### Behavioral Assessments

Motor performance was assessed in a speed test, termed the “30-second challenge”. Here, sequence accuracy was calculated as the number of successfully executed repetitions of a sequence during a 30-s period (Karni et al., 1995). Participants were presented with four visual images of sequences identical to their trained sequences, and four untrained sequences (which were the reverse of trained sequences to match the difficulty). In this test, the sequences were displayed as a static image, presented in counter-balanced order, and finger-to-thumb touches data were recorded using an adapted Peregrine Gaming Glove (https://peregrineglove.com/), with a software designed by Minerva Design, UK. Participants were also filmed by video camera using a Canon PowerShot SX240 HS for verification.

### Procedure

Participants underwent an initial MRI and behavioral assessment prior to the four-week training (pre-training assessment). Next, participants trained at home three times per week for 20 minutes per session by watching four five-minute training videos online and logging their progress which was monitored by the researchers. Each video included a slow increase in tempo, and four levels with increasing starting speed were provided, so that participants could opt to increase the speed of the visual display and tempo of the music, if the previous level was no longer challenging.

Each video ended with the “30-s challenge”, which served as the behavioral assessment in each time point, and none of the participants reached the final level indicating sufficient difficulty for learning progression. Participants came into the lab for a mid-training behavioral assessment, two weeks after the start of the training (mid-point assessment), and completed both behavioral testing and MRI scanning after the four-week training concluded (post-training assessment).

### MRI Data Acquisition

Participants were scanned before and after the training at the Clinical Research Imaging Centre (CRIC), University of Edinburgh, UK (http://www.cric.ed.ac.uk), using a 3 Tesla Siemens Verio system (Siemens Healthcare, Erlangen, Germany) equipped with a 12-channel matrix head coil. T1-weighted images were acquired using an MPRAGE gradient-recalled-echo (GRE) sequence (voxel size: 1 mm isotropic; 176 slices; field of view (FOV): 256 × 240 × 176 mm; repetition time [TR]: 2.3 ms; echo time [TE]: 2.98 ms; inversion time [TI]: 900 ms; flip angle: 9°), with a bandwidth of 238 Hz/pixel. Partial Fourier sampling (factor = 1) and parallel imaging (GRAPPA

acceleration factor = 2) were applied to optimize scan efficiency. Diffusion-weighted images were acquired using a single-shot spin-echo echo-planar imaging (EPI) sequence (b = 1000 s/mm^2^; 56 non-collinear directions), along with six T2-weighted (b = 0 s/mm^2^) volumes. Scans were obtained in the axial plane ([FOV]: 240 × 240 mm; 55 contiguous slices; voxel size: 2.5 mm isotropic). Acquisition parameters were: [TR] = 10.2 s; [TE] = 69.6 ms; bandwidth = 2126 Hz/pixel.

### Preprocessing of MRI Data

The longitudinal pipeline of FreeSurfer (v.7.4.1, http://surfer.nmr.mgh.harvard.edu; Reuter et al., 2012a) was used to preprocess the T1-weighted images. Data were first processed cross-sectionally, after which a subject-specific template (Reuter & Fischl, 2011) was created by registering and averaging both time points to be transformed into the template space to reduce within-subject variability and to be able to detect longitudinal changes over time (Reuter et al., 2010). Pre-processing of this data consisted of normalization, Talairach transformation, removal of non-brain tissue, intensity correction, as well as cortical parcellation (Reuter et al., 2012b).

Cortical thickness was computed as the distance between the grey-white matter boundary and the pial surface at each vertex of the tessellated cortical surface (Dale et al., 1999; Fischl & Dale, 2000). Grey matter volume, and surface area were derived from the reconstructed surfaces using segmentation and parcellation in FreeSurfer. These metrics were then extracted for each participant using the Desikan-Killiany atlas (Desikan et al., 2006). Diffusion tensor imaging data was preprocessed using FSL Tools (http://www.fmrib.ox.ac.uk/fsl) for brain extraction, correction for head motion, and eddy current distortions by registering all volumes to the first

T2-weighted image (Jenkinson & Smith, 2001). The diffusion tensor was estimated using DTIFIT, and voxel-wise maps of FA were derived from the tensor eigen values. Tract-average values of FA within the bilateral arcuate fasciculi were obtained using probabilistic neighborhood tractography (Clayden et al., 2011). The TractoR package (http://www.tractor-mri.org.uk), was used to create models of the bilateral arcuate fasciculi, which were segmented for each subject from underlying connectivity data produced using FSL’s BedpostX/ProbTrackX algorithm (Behrens et al., 2003). Three participants were removed from the DTI analysis after the tract segmentation was reviewed by an experienced and blinded operator, who deemed the tracts anatomically implausible (for a full description, see Moore et al., 2017).

#### Regions of interest (ROIs)

ROIs were defined using the Desikan-Killiany atlas (Desikan et al., 2006), covering the abovementioned functional regions, implemented as two parcellations covering auditory regions, namely the Superior Temporal Gyrus (STG), and Transverse Temporal Gyrus (TTG); as well as the following motor regions: Precentral Gyrus (PCG), and the Paracentral Lobule (PCL; see Figure 1).

**Figure 1.**
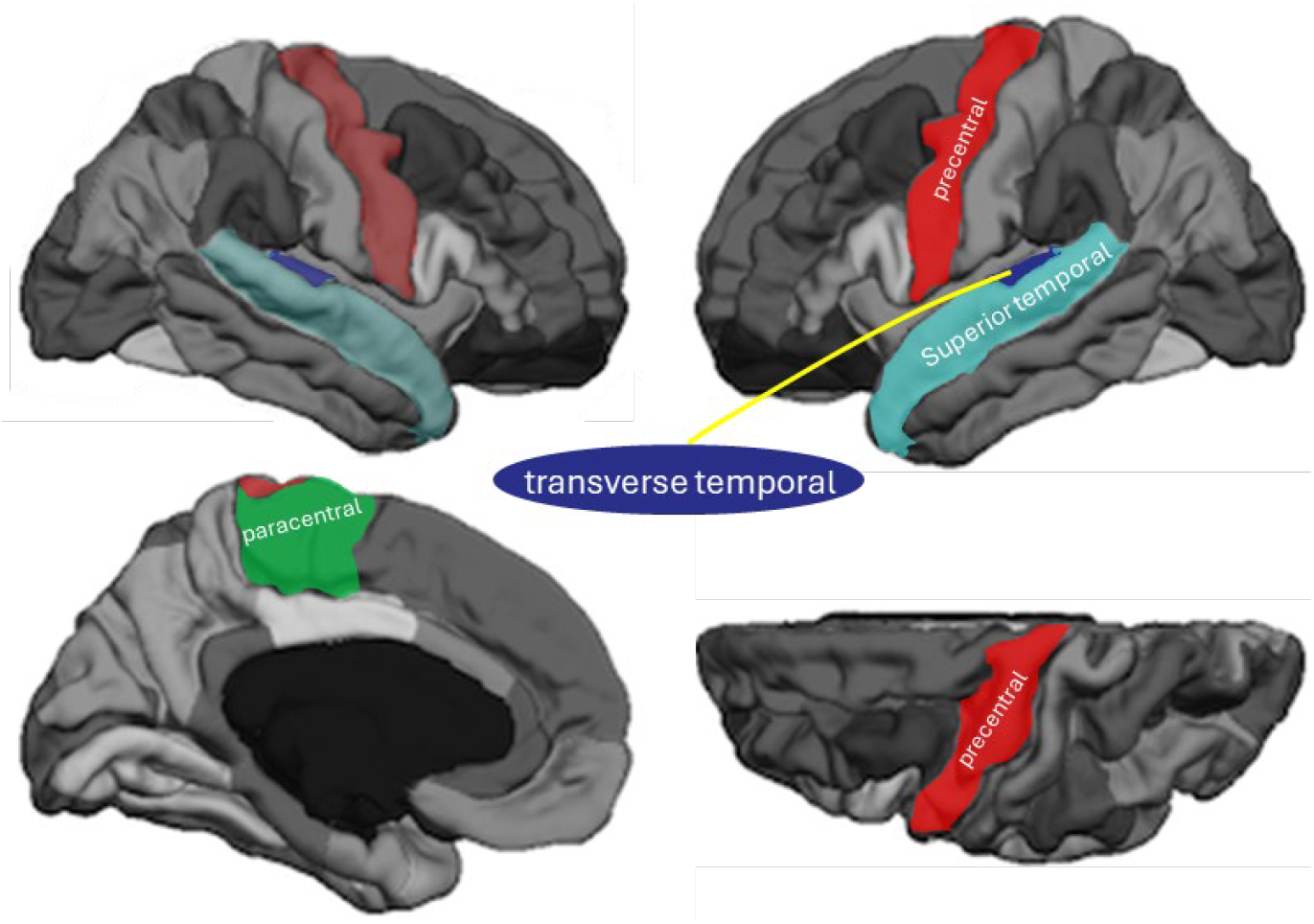
Parcellation and segmentation based on the Desikan-Killiany atlas in FreeSurfer. *Note:* Regions of interest are highlighted, specifically the auditory (superior and transverse temporal gyrus) and motor (precentral gyrus and paracentral lobule) areas. Image is based on Klein & Tourville, 2012.

### Statistical Analysis

#### Behavioral Analysis

The number of correctly performed sequences for each trial was extracted and averaged for the Trained and Untrained sequences pre-, mid-, and post-training using MATLAB 2010a (Natick, USA). The data were checked for normality of distribution, both by visually inspecting histogram plots, and using the Shapiro-Wilk’s test with a *p-*value threshold of 0.05. Homogeneity of variance was checked using Levene’s test with a significance *p-*value of 0.05. Outliers were identified using three multiples of the interquartile range.

Sequence accuracy for each of four sequences was averaged, pre, mid, and post-training separately for trained and untrained sequences in R (R Core Team, 2024). Performance on each measure for trained and untrained sequences was compared across the three time points and between groups (Music *vs*. Control) in a mixed ANOVA, using the R package “ez” (Bakeman, 2005; Lawrence, 2016). Bonferroni correction of the α level was applied to account for multiple comparisons.

#### Grey Matter Analysis

Cortical measures of thickness, volume, and surface area were extracted for analysis in R. The “brms” R package (Bürkner, 2017), which interfaces with “rstan” for Bayesian inference (Bürkner, 2021). Inference was based on 90% credible intervals (CrI), which is considered the default in Bayesian models. Moreover, 90% CrIs are more stable compared to 95% interval, which would require a substantially large sample size > 10,000 (Makowski, Ben-Shachar, & Lüdecke, 2019). Bayesian models do not rely on a *p*-value with a cut-off threshold to determine significance (Kruschke, 2013), rather if CrIs do not overlap 0, the effect is considered to have 90% probability of falling within that range. In case 0 was slightly overlapped, probability of direction (*Pd*), as calculated by the R package “bayestestR” (Makowski, Ben-Shachar, & Lüdecke, 2019), was used to determine the probability of an effect being strictly positive or negative, and is an index of effect existence (Makowski, Ben-Shachar, Chen, et al., 2019).

Bayesian linear mixed effects models were conducted for each of the three outcome measures (thickness, volume, surface area) of each ROI (STG, TTG, PCG, PCL), which were Z-score transformed such that 0 is the mean, and with a standard deviation of 1 (Barreda & Silbert, 2023). Fixed effects (predictors) were modeled as binaries, specifically Groups (Music vs. Control), hemisphere (Right vs. Left), time-point (Pre vs. Post training) and their two-way and three-way interactions were modeled to test for effects of training on brain structure. All models assumed a Gaussian likelihood (Bürkner, 2018). Participants were added as the random effect, with varying intercepts and slopes for time-points, and hemispheres to account for repeated measures, and individual variability. The hypothesis function was applied to assess if the main effects of interest (Time × Group, and Time × Group × Hemisphere) were credibly different from zero, and if evident, were followed up to probe the source of the interaction.

Convergence was assessed based on Rhat 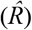 statistic, effective sample sizes, and visual inspection of trace plots. All models demonstrated excellent convergence, with 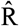 values below 1.01 and effective sample sizes > 1,000 for all parameters, indicating stable and reliable posterior

estimates. Posterior predictive checks revealed a good model fit for all assessed models (Makowski, Ben-Shachar, & Lüdecke, 2019). For full reporting of model parameters and outcomes, according to reporting guidelines (Keysers et al., 2020; Kruschke, 2021), and studies reporting on similar methods (Erdman et al., 2025; Worschech et al., 2022) see Supplementary Material S1.

#### Brain-Behavior Relationships

The associations between the change pre-to-post training in sequence accuracy and the change in grey matter cortical measures (thickness, volume, area) for each ROI (STG, TTG, PCG, PCL) over all participants, and for each group separately, was examined using Spearman (rather than Pearson) correlations to take inhomogeneity of variances into account. To address normality violations, a Yeo-Johnson transformation was applied by automatically estimating the best lambda value for normalization using the “bestNormalize” R package (Peterson & and Cavanaugh, 2020), the Yeo-Johnson transformed values were used for all correlational analyses. Bonferroni correction of the α level is applied to correct for multiple comparisons.

#### Grey and White Matter Relationships

Finally, as the choice of ROIs examined in the cortical grey matter analysis was motivated by our previous findings on white matter connectivity (Moore et a., 2017), we also examined Spearman correlations between fractional anisotropy (FA) of the arcuate fasciculus and cortical measures (thickness, volume, and area) of the ROIs for each group, and within hemispheres. An identical procedure of the correlational analyses above was followed. Three participants were removed from the DTI analysis, as their tract segmentation was deemed anatomically implausible.

## Results

### Behavioral Change in Sequence Accuracy Over Time

Descriptives of the behavioral measures are shown in Table 1. A repeated-measures ANOVA examining motor sequence accuracy revealed significant differences between time points [*F*(2, 56) = 33.90, *SE* = 69.01, *p* <.001, *η*^2^*g* = 0.11], trained *vs*. untrained sequences [*F*(1, 28) = 28.09, *SE* <0.00, *p* <.001, *η*^2^*g* = 0.026)] and their interaction [*F*(2, 56) = 16.55, *SE* = 15.63, *p* <.001, *η*^2^*g* = 0.013], while no group differences or group interactions were found (*p* >.05). Post-hoc pairwise comparisons revealed significant differences between all time-points for the trained sequences, and between the first two time points of the untrained sequences (indicating that only for trained sequences, performance significantly increased in the second half of the training), with all changes showing increases in sequence accuracy over time (*p* <.001, *d* = 0.33 – 1.02). Finally, participants also had better sequence accuracy on the Trained > Untrained Sequences, at mid-(*p* <.001) and post-training (*p* <.0001) but not at pre-training (*p* >.05), indicating sequence-specific learning. For a visual representation, see Figure 2.

**Table 1.**
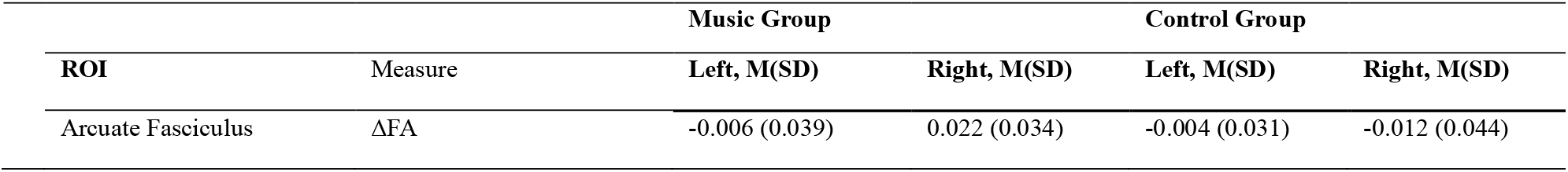
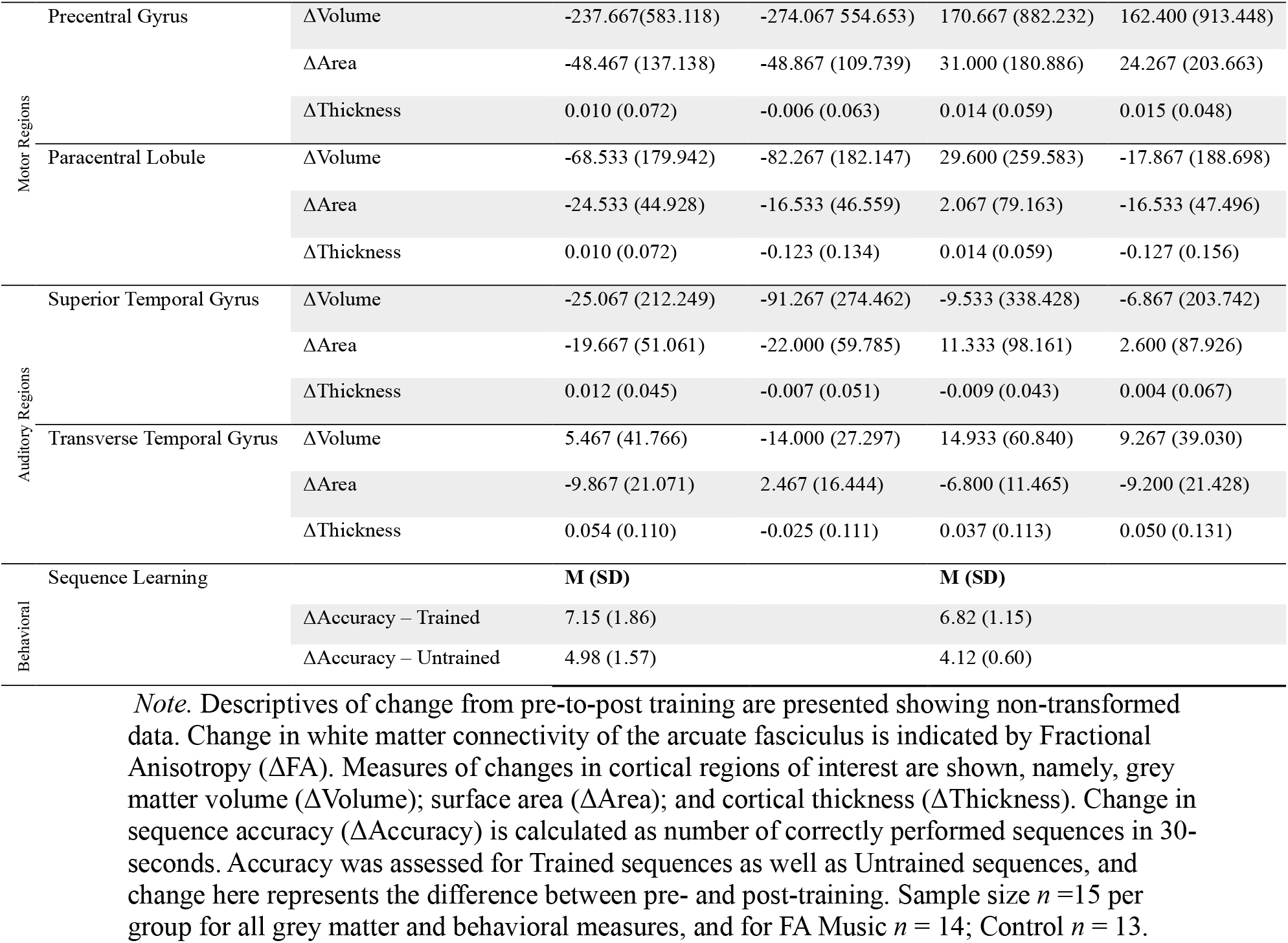
Means and standard deviations of change in brain measures pre- to post-training per group and hemisphere in all ROIs, as well as the change in behavioral sequence accuracy.

**Figure 2.**
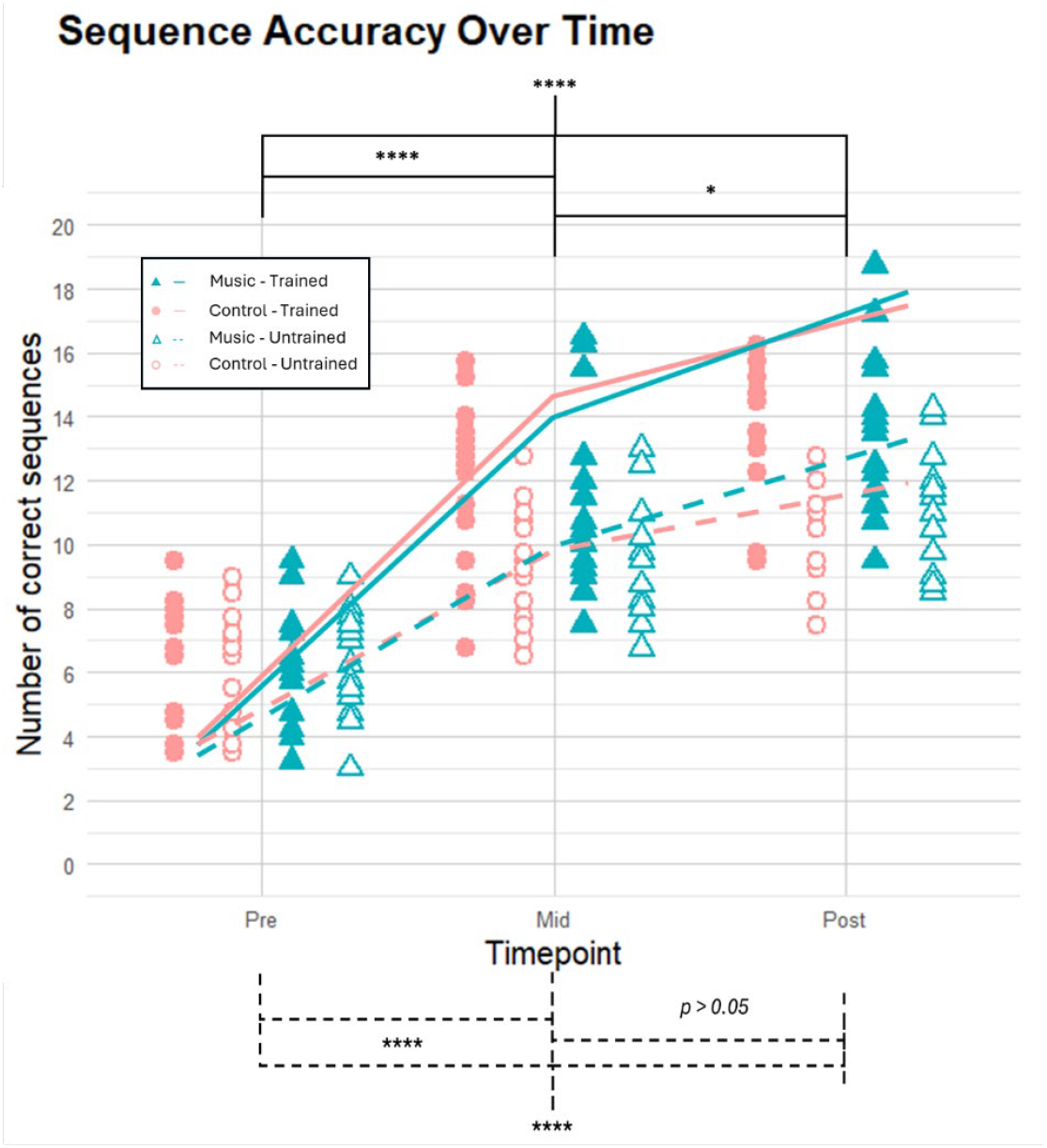
Sequence accuracy by group and sequence type at three timepoints. *Note*. Sequence accuracy is displayed across three time points, with trained sequences shown in solid lines/colored data points and untrained sequences in dashed lines/hollow data points. Different colors indicate group conditions (Music vs. Control). Reported *p*-values are bonferroni adjusted. ^(****)^ indicates significance is below <.0001-level, and (*) signficance below <.05-level, shown for the trained sequences on the top and untrained sequences on the bottom.

### Grey Matter Changes Pre-and-Post Training, and Group-Hemisphere Interactions

Change (post-minus pre-training) values for all ROIs for each hemisphere and group are presented in Table 1, alongside change values in behavioral performance and FA in arcuate fasciculus.

Bayesian linear mixed effects models were estimated for all z-score transformed grey matter outcomes to assess the effects of hemisphere, timepoint, group and their interactions on cortical structure measures (see Supplementary S1 for full model outcomes). The analysis revealed moderate evidence (*β* = 0.17, 90% CrI [0.00, 0.35], *Pd* = 95.10%) for a three-way interaction of Time × Group × Hemisphere in the model predicting the surface area of the TTG. The three way interaction suggests a small but credible effect of training on TTG surface area, showing greater increase in the Music group in the right hemisphere over time compared to the control (*β* = 0.14, [0.01, 0.27]), whereas the Control showed a decrease in right hemisphere over time (-0.11, [-0.20, -.01]). In the left hemisphere TTG’s surface area decreased in the Music group (-0.12, [-0.21, -0.02]), but this was not evident for the Control (-0.08, [-0.18, 0.02]), nor was the change different between groups (-0.04, [-0.17, 0.10]), see Figure 3 for the main outcome, as well as Figure S1 and Table S4 in Supplementary Material S2 for detailed results.

**Figure 3.**
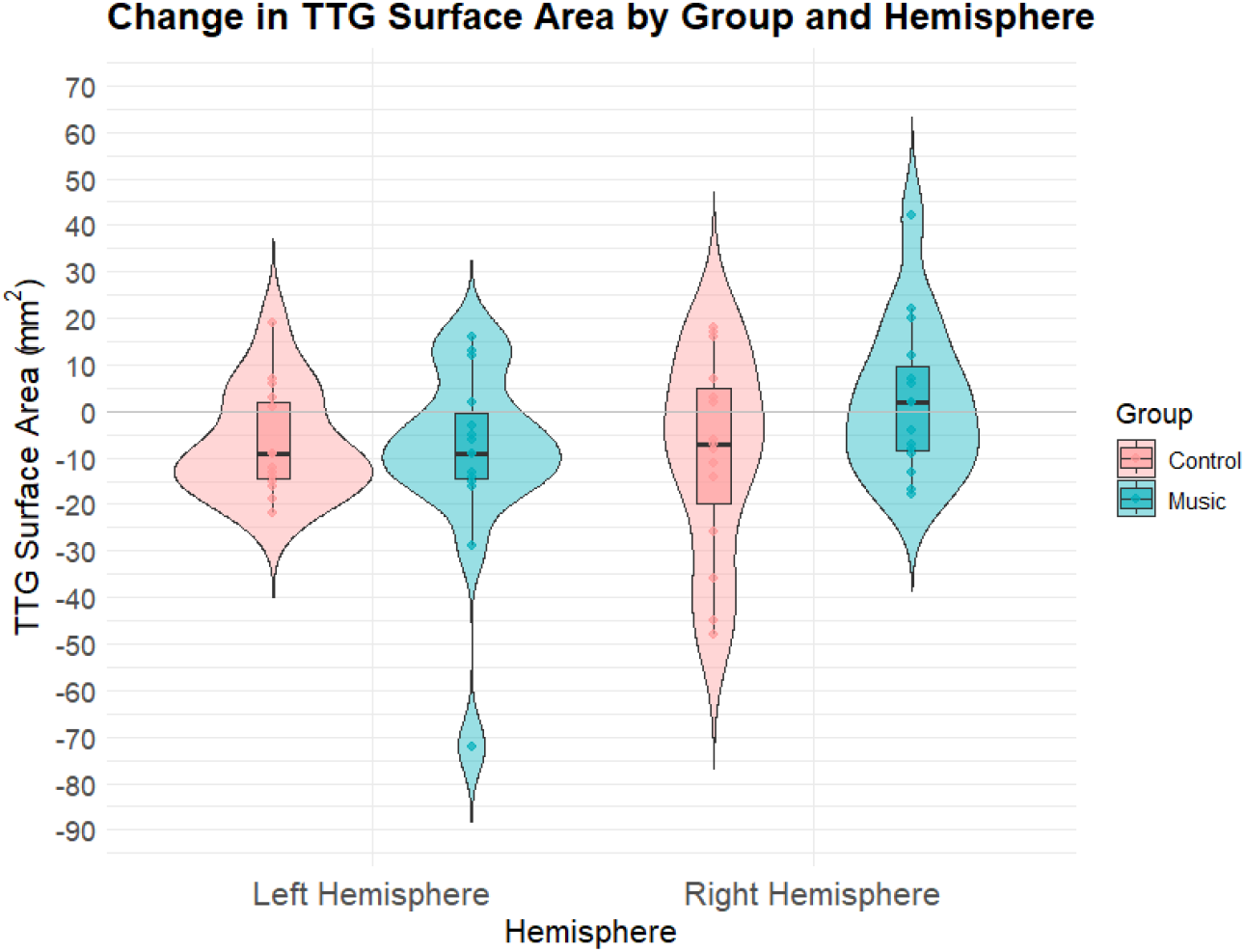
Change in Surface Area of TTG between Groups and Hemispheres. *Note*. Change in surface area of the transverse temporal gyrus (ΔTTG Area) is shown for both hemispheres (LH on the left and RH on the right) using non-transformed data for both groups, shown in different colors.

Although the model of STG cortical thickness revealed moderate evidence of a three-way interaction (*β* = -0.28, [-0.53, -0.03], *Pd* = 96.6%), there were no differences over time between or within groups. Except for the three-way interaction itself, the credible intervals of these effects all overlap zero when examining their probability of direction, indicating that these effects are not meaningful, for more details see Figure S2 and Table S5 in Supplementary Material S2.

No other models showed evidence for changes over time, or interactions of Time with × Group, Time × Hemisphere, or a three-way interaction. Posterior probabilities for these terms remained below conventional thresholds of 95% (also see Supplementary Material S1). While a number of findings emerged that did not interact with time (time-group or time-group-hemisphere), these cannot be attributed to the training, and are thus not relevant to our hypotheses.

Figure 3 displays the interaction effect between hemisphere and group for the change of surface area over time of the TTG, showing that for the music group, the right TTG increased while the left TTG decreased, while the Control group showed decreases in both hemispheres. Of these changes, only the difference between groups in the right hemisphere, the decrease over time in the left hemisphere of the Music Group, and decrease over time in the right hemisphere of the Control group, as well as the difference between the hemispheres in the Music group showed meaningful change over time, evident by CrI values that do not include 0 and *Pd* > 95%.

### Brain-Behavior Relationships: Change in Motor Learning and Cortical Structures

When examining associations between change in sequence accuracy and change in cortical measures of ROIs, several negative moderate associations were found in the Music but not in the Control group, however, none of these associations survived multiple comparison correction (*Bonferroni corrected* ***α*** = 0.004). Specifically, in the Music (but not in in the Control Group) PCG surface area change of the left hemisphere was negatively associated with performance change in both trained and untrained sequences (*ρ =* -0.53 – -0.58; *p* = 0.02 – 0.04, -*pBonferroni* =.24 –.48), the latter was also found when examining all participants from both groups together. Whereas only in the Music Group, the bilateral PCL surface area change was negatively associated with trained sequences change (*ρ =* -0.52 – -0.54, *p* = 0.03 – 04, *p-Bonferroni* =.36 –.48). For more detailed results, see Supplementary Materials S3.

### Grey and White Matter Relationships

Descriptives of the FA for the arcuate fasciculus, as reported in Moore et al. (2017), are included in Table 1. Post-hoc correlations between changes in grey matter measures (thickness, volume, and area) of the ROIs and changes in FA of the arcuate fasciculus revealed some group-specific lateralized associations. As three participants were removed from this analysis, the participants included 14 in the Music Group (*M*age = 21.36, *SD* = 2.02 years; 3 males) and 13 in the Control (*M*age = 21.54, *SD* = 2.47 years; 4 males) group. Although Spearman’s *ρ* indicated moderate to strong associations, none of these survived Bonferroni corrections (*Bonferroni corrected* ***α*** = 0.004). The Music group showed an inverse relationship between FA change of the left arcuate and volume change of the left PCG (*ρ* = -0.604, *p* =.025, *p-Bonferroni* =.30). Conversely, the Control group showed positive associations in the right hemisphere between FA change of the arcuate fasciculus and volume change of the PCL, and surface area change of the PCG (*ρ* =.61 –.73, *p* =.01 –.03, *p-Bonferroni* =.12 –.36). For detailed results, see Supplementary Materials S4.

## Discussion

The current study examined changes in motor performance related to auditory and motor cortical regions over a 4-week visuomotor training, using the left hand, with or without musical cues.

The results of this study align with previous findings of motor learning and white matter change on the same dataset (Moore et al., 2017). The addition of musical cues to a visuomotor sequence training task was expected to lead to changes in cortical thickness (and by association volume and surface area) of the auditory cortex (STG and TTG), with similar changes expected in the primary and premotor regions (PCG and PCL) of the right hemisphere contralateral to the trained hand. The study further explored how behavioral changes in sequence accuracy and change in FA of the arcuate fasciculus correlated to the cortical measures of these regions.

Behaviorally, there were no differences between groups in motor performances. However, learning was seen over time over all participants, clearly showing better performance on trained than untrained sequences, indicating both general as well as sequence-specific motor learning. Examining the change from pre-to post-training in grey matter indices in auditory and motor brain regions between groups and hemispheres revealed moderate evidence of a time-group-hemisphere interaction for the surface area of the TTG (or Heschl’s gyrus), showing that the Music group showed an increase in surface area of the right TTG, while the Control group showed a decrease (Figure 3). While a three-way interaction was similarly evident for the STG’s cortical thickness, none of the hypothesized comparisons with regards to change over time within or between groups yielded evident differences. No other differences related to change over time, between groups, hemispheres, or three-way interactions were found for any cortical indices of the examined auditory or motor regions. Although associations between improvements on sequence accuracy and change in cortical measures of our regions of interest were seen only in the Music group, these did not survive correction for multiple comparisons. Similarly, when exploring associations between the change in cortical grey matter measures and previously reported increased connectivity of the arcuate fasciculus, some group-specific lateralized associations were observed, but none survived multiple comparison correction. The increase of surface area only in the right Heschl’s gyrus of the Music group highlights the difference between training with or without musical cues, revealing changes contralateral to the trained in the auditory cortex only with music-cued training. However, several expected findings did not emerge. While the expected right hemispheric involvement was observed for the auditory cortex, it was not evident for motor regions. Below, we discuss plausible interpretations of the current findings in relation to previous research on music-based motor learning.

### Interhemispheric changes between groups over time

Concerning the primary research question, the hypotheses were only partly confirmed. Between-group differences over the four-week training were seen in the auditory cortex, where the Music group had an increase in surface area of the right and a decrease in the left hemisphere, whereas the Control group showed a decrease in both (Figure 3), reflecting the same pattern as a previous finding in older adults following a 6-month piano training (Worschech et al., 2022). The increase in surface area of the right Heschl’s gyrus in the Music Group may reflect increased auditory-motor coupling based on music-cued motor learning in the Music group, in line with previously found increase in FA in the right hemisphere, contralateral to the trained hand (Moore et al., 2017). The fact that this was only evident in the right hemisphere may indicate auditory-motor connectivity relating to the nature of the trained task involving the left hand (contralateral to the right hemisphere finding). Additionally, previous findings suggest that pitch, meter, and pattern processing lateralize to the right temporal lobe (Peretz & Zatorre, 2005; Thaut et al., 2014), and working memory involvement in melody learning is also thought to be right lateralized (Peretz & Zatorre, 2005), which may further underlie the current finding. However, no effects in cortical volume or thickness were seen within or between groups over time. Moreover, our hypotheses on changes in motor areas were not supported. The absence of any changes in these areas, as termination points of the arcuate fasciculus, may be attributed to the fact that this previous finding on white matter connectivity is interpreted as related to the connection between auditory and motor regions contralateral to the trained hand (Moore et al., 2017), while in this study these structures are examined separately. Although other studies reported changes in cortical motor areas after learning a finger-to-thumb movement sequence (Sale et al., 2017), the current investigation did not lead to measurable change in motor regions, and adding musical cueing did not affect learning-related plasticity overall.

### Relationships between changes in behavioral and cortical measures

Although not surviving multiple comparison corrections, medium-sized associations were found only for the Music group between increases in sequence accuracy and decreases in surface area of bilateral and ipsilateral cortical motor areas (detailed in S3). These results cautiously suggest that changes in these motor regions may thus be more related to performance increases specifically when learning a movement with musical cueing, even though the direction is unexpected. While increases in surface area and decreases in cortical thickness have been related to increased white matter connectivity (Storsve et al., 2014), this refers to relations between neuroimaging modalities rather than the relationship with behavioral measures of learning reported here. However, the finding that these mechanisms might be affected through musical cues is new, and given the tenuousness of these findings, should be further explored in future research.

### Relationship between cortical grey matter changes and white matter changes

To align with the previous finding that music-cued training led to increased white matter connectivity in the right arcuate fasciculus, contralateral to the trained hand (Moore et al., 2017), associations between this change in FA and grey matter indices in auditory and motor regions were assessed post-hoc. Again, several associations were identified that did not survive multiple comparison corrections (detailed in S4). Here, unexpected dissociations of group and hemisphere suggest differential associations between grey and white matter structures that distinguish motor learning with and without musical cues. However, as these results are not evident after multiple comparison correction, and interactions between FA values and grey matter indices are likely mostly physiological rather than based on cueing of motor training, this is not interpreted further.

### Limitations & Future Directions

This study had a modest sample size of 15 participants per group. To substantiate these findings, future studies should seek to include a larger sample size, which is especially relevant for the detection of small-to-medium sized neural effects reported in this study. Additionally, this sample of young, non-musician participants may not generalize to other groups where potential for plasticity may be different, such as in older adults or people suffering from neurological disorders. Furthermore, this paradigm only utilizes finger movements and as such focuses only on fine motor skills. More generally, it has been questioned whether structural changes measured in MRI are reflective of true grey matter changes, or a particularity of the imaging modality capturing vascular or other alterations, as opposed to true neuronal plasticity (Garzón et al., 2023). The current study, showing mixed outcomes relating grey and white matter measures, does not provide a more robust account of the relations between imaging modalities. Moreover, there may be different results in other ROIs that are involved that have not been included here, for instance the inferior frontal gyrus, which is also related to the arcuate fasciculus, but arguably has a less primary function in music-cued movement than motor or auditory areas. Finally, the way behavioral learning was assessed did not focus on temporal accuracy, but rather indexes learning based on self-timed movements performed as fast as possible. Importantly, while visual cues are shown to support learning motor sequences, auditory cues seem to be crucial for motor timing (Comstock et al., 2018; Mate, 2022). Future studies should preferably measure timing accuracy in the same context as it was trained, instead of in silence using still images.

With health-related applications in mind, future efforts should include populations that are more likely to need health interventions based on cued movement. Previously, grey matter changes have been demonstrated after six months of piano training compared to no motor training in elderly participants (Worschech et al., 2022), and even though that study did not investigate the effect of adding musical cues to an identical motor training, it does confirm the potential for plasticity in this group. The current study not only demonstrates that it is possible to observe changes in grey matter structure in the span of weeks in young adults, but specifically highlights that adding musical cues leads to distinct neuroplastic changes.

### Conclusions and Implications

This study demonstrates that adding musical cues to a short-term motor training induces changes in right primary auditory cortex in a sample of healthy, young-adult non-musicians after four weeks of left-hand movement training, specifically showing increased surface are of the right Heschl’s gyrus after only 12 sessions of music-cued practice in an experimentally controlled study. While there are some indications that adding musical cues impacts on specialization in motor areas related to behavioral learning, this needs to be further investigated. These findings are the first to show grey matter changes related to musical cues in motor learning, and are in line with our previous findings on white matter changes in the same context. This provides a basis for future work addressing the potential for plasticity and health-based applications using musical cueing in more diverse populations.

## Authors Contributions

Author contributions: R.S., N.R. and K.O. designed the study, R.S. and K.O. created the training paradigm. E.M. collected the data. Analysis plan was devised by M.M., D.V., S.R., K.O. and R.S., analysis for behavioral, and MRI data was performed by M.M. D-MRI data were analyzed by E.M. and M.B. The first draft of the paper was provided by M.M with comments and edits by R.S. and K.O., and all authors contributed edits and approved the final version.

## Acknowledgments

The authors would like to thank Elise S. Besemer for her assistance in preprocessing the MRI data, Dr. Franz Wurm for his input on reporting of Bayesian statistics, and Dr. Zack Moir for composing the music used in the training videos.

