## Supplementary Materials for "Increased surface area of Heschl’s gyrus following music-cued motor training"

### Supplementary Materials S1:

Table S1, S2, and S3 show the model outcomes of the Bayesian multilevel models on each of the cortical measures (cortical thickness, grey matter volume, and surface area) for each of the ROIs (namely STG, TTG, PCG, and PCL). Markov chain Monte Carlo (MCMC) with four chains, each with iter = 8,000, warmup = 4,000, thin = 1; total post-warmup samples = 16,000; Adapt_delta = 0.80; max_treedepth = 10; family = Gaussian; backend = “rstan”; application of weakly informative priors for fixed effects, random intercepts, and residual SD.

Table S1. Model outcomes for the Bayesian multilevel models on cortical thickness.

| **Model Formula:** Zscore ~ Time * Group * Hemisphere + (1 + Time + Hemisphere\| Participant) | | | | |
|---|---|---|---|---|
| Measure: Cortical Thickness | **STG** | **TTG** | **PCG** | **PCL** |
| **Group-Level Effects** | | | | |
| sd(Intercept) | **1.04 [0.79, 1.40]** | **0.95 [0.70, 1.29]** | **0.89 [0.67, 1.19]** | **0.93 [0.69, 1.25]** |
| sd(Time) | **0.39 [0.24, 0.57]** | **0.20 [0.01, 0.48]** | **0.42 [0.28, 0.60]** | **0.49 [0.32, 0.70]** |
| sd(Hemisphere) | **0.75 [0.55, 1.01]** | **0.67 [0.42, 0.97]** | **0.95 [0.72, 1.27]** | **0.84 [0.62, 1.14]** |
| cor(Intercept,time) | 0.07 [-0.34, 0.49] | 0.04 [-0.72, 0.82] | -0.13 [-0.50, 0.30] | -0.10 [-0.49, 0.32] |
| cor(Intercept,Hemisphere) | -0.36 [-0.66, 0.01] | -0.21 [-0.59, 0.25] | -0.27 [-0.59, 0.11] | -0.28 [-0.60, 0.11] |
| cor(Time,Hemisphere) | -0.19 [-0.59, 0.25] | 0.06 [-0.78, 0.82] | 0.31 [-0.09, 0.65] | 0.31 [-0.11, 0.66] |
| **Population-Level Effects** | | | | |
| Intercept | -0.27 [-0.82, 0.27] | -0.32 [-0.85, 0.21] | -0.01 [-0.49, 0.46] | -0.34 [-0.84, 0.15] |
| Time | -0.08 [-0.34, 0.17] | 0.19 [-0.14, 0.50] | 0.11 [-0.15, 0.38] | 0.10 [-0.21, 0.41] |
| Group(MG) | 0.22 [-0.57, 1.00] | 0.34 [-0.41, 1.07] | 0.23 [-0.45, 0.90] | 0.40 [-0.27, 1.10] |
| Hemisphere(Right) | 0.30 [-0.13, 0.70] | 0.19 [-0.26, 0.65] | **-0.51 [-1.02, 0.01]** | 0.24 [-0.23, 0.71] |
| Time:Group(MG) | 0.19 [-0.18, 0.55] | 0.08 [-0.35, 0.53] | -0.03 [-0.41, 0.34] | -0.18 [-0.61, 0.26] |
| Time:Hemisphere(Right) | 0.11 [-0.11, 0.33] | 0.06 [-0.34, 0.47] | 0.01 [-0.21, 0.21] | 0.06 [-0.19, 0.31] |
| Group(MG):Hemisphere(Right) | 0.07 [-0.51, 0.65] | -0.06 [-0.70, 0.58] | 0.46 [-0.28, 1.17] | 0.01 [-0.65, 0.65] |
| Time:Group(MG):Hemisphere(Right) | **-0.28 [-0.58, 0.02]** | -0.45 [-1.02, 0.12] | -0.14 [-0.43, 0.16] | 0.00 [-0.36, 0.37] |
| **Family-Specific Parameters** | | | | |
| sigma | -0.28 [0.53, -0.03] | 0.40 [0.30, 0.50] | 0.21 [0.16, 0.28] | 0.25 [0.19, 0.34] |
| **Posterior predictive check** | 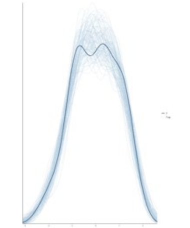 | 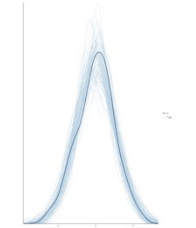 | 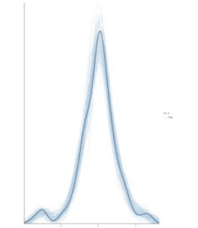 | 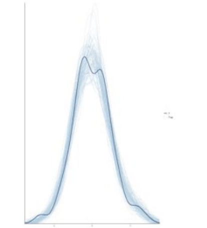 |

*Note.* The table shows group level, and population level effects, the z-score transformed values of the median estimate is followed by the credible intervals between brackets. The columns present each of the ROIs, namely, STG: Superior Temporal Gyrus; TTG: Transverse Temporal Gyrus; PCG: Precentral Gyrus; and PCL: Paracentral Lobule. SD indicates standard deviation, Cor indicates correlation between parameters with varying slopes. MG: Music Group. Effects that are pertinent to the hypotheses are in black, whereas all other effects are faded in grey. Effects that are credibly different from zero are in bold setting.

Table S2. Model outcomes for the Bayesian multilevel models on grey matter volume.

| **Model Formula:** Zscore ~ Time * Group * Hemisphere + (1 + Time + Hemisphere\| Participant) | | | | |
|---|---|---|---|---|
| Measure: Grey Matter Volume | **STG** | **TTG** | **PCG** | **PCL** |
| **Group-Level Effects** | | | | |
| sd(Intercept) | **1.17 [0.90, 1.54]** | **0.85 [0.65, 1.12]** | **1.13 [0.87, 1.51]** | **0.87 [0.66, 1.15]** |
| sd(Time) | **0.14 [0.07, 0.20]** | **0.06 [0.00, 0.14]** | **0.52 [0.39, 0.69]** | **0.39 [0.27, 0.53]** |
| sd(Hemisphere) | **0.87 [0.66, 1.16]** | **0.76 [0.58, 1.00]** | **0.86 [0.65, 1.15]** | **1.21 [0.92, 1.60]** |
| cor(Intercept,time) | -0.22 [-0.60, 0.21] | -0.38 [-0.92, 0.57] | -0.41 [-0.68, -0.06] | -0.30 [-0.62, 0.08] |
| cor(Intercept,Hemisphere) | **-0.50 [-0.74, -0.19]** | **-0.62 [-0.82, -0.35]** | -0.35 [-0.64, 0.00] | -0.30 [-0.61, 0.06] |
| cor(Time,Hemisphere) | 0.08 [-0.35, 0.50] | 0.17 [-0.67, 0.84] | 0.19 [-0.18, 0.53] | -0.17 [-0.53, 0.22] |
| **Population-Level Effects** | | | | |
| Intercept | 0.25 [-0.35, 0.84] | **0.80 [0.36, 1.23]** | 0.03 [-0.55, 0.61] | **-0.55 [-1.00, -0.11]** |
| Time | -0.01 [-0.10, 0.09] | 0.07 [-0.04, 0.17] | 0.11 [-0.16, 0.39] | 0.06 [-0.17, 0.29] |
| Group(MG) | -0.14 [-0.98, 0.70] | -0.24 [-0.88, 0.38] | 0.15 [-0.68, 0.97] | **0.66 [0.01, 1.31]** |
| Hemisphere(Right) | **-0.42 [-0.88, 0.04]** | **-1.49 [-1.89, -1.09]** | -0.17 [-0.62, 0.28] | **0.61 [-0.02, 1.25]** |
| Time:Group(MG) | -0.01 [-0.14, 0.12] | -0.04 [-0.19, 0.10] | -0.27 [-0.66, 0.12] | -0.20 [-0.52, 0.12] |
| Time:Hemisphere(Right) | 0.00 [-0.08, 0.09] | -0.02 [-0.16, 0.11] | -0.01 [-0.11, 0.10] | -0.09 [-0.24, 0.06] |
| Group(MG):Hemisphere(Right) | 0.17 [-0.48, 0.80] | 0.23 [-0.33, 0.81] | -0.02 [-0.66, 0.62] | -0.21 [-1.13, 0.69] |
| Time:Group(MG):Hemisphere(Right) | -0.04 [-0.17, 0.08] | -0.06 [-0.25, 0.14] | -0.02 [-0.17, 0.13] | 0.07 [-0.15, 0.28] |
| **Family-Specific Parameters** | | | | |
| sigma | 0.08 [0.06, 0.12] | 0.13 [0.11, 0.17] | 0.10 [0.08, 0.13] | 0.14 [0.11, 0.19] |
| **Posterior predictive check** | 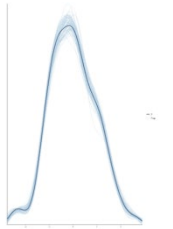 | 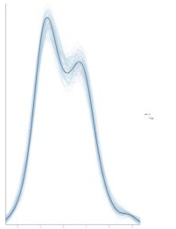 | 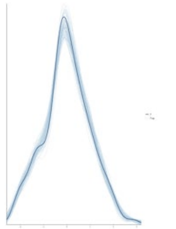 | 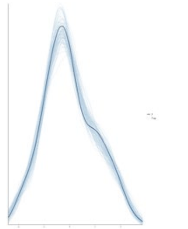 |

*Note.* The table shows group level, and population level effects, the z-score transformed values of the median estimate is followed by the credible intervals between brackets. The columns present each of the ROIs, namely, STG: Superior Temporal Gyrus; TTG: Transverse Temporal Gyrus; PCG: Precentral Gyrus; and PCL: Paracentral Lobule. SD indicates standard deviation, Cor indicates correlation between parameters with varying slopes. MG: Music Group. Effects that are pertinent to the hypotheses are in black, whereas all other effects are faded in grey. Effects that are credibly different from zero are in bold setting.

Table S3. Model outcomes for the Bayesian multilevel models on surface area.

| **Model Formula:** Zscore ~ Time * Group * Hemisphere + (1 + Time + Hemisphere\| Participant) | | | | |
|---|---|---|---|---|
| Measure: Surface Area | **STG** | **TTG** | **PCG** | **PCL** |
| **Group-Level Effects** | | | | |
| sd(Intercept) | **1.13 [0.87, 1.49]** | **0.80 [0.61, 1.06]** | **1.12 [0.86, 1.49]** | **0.90 [0.68, 1.21]** |
| sd(Time) | **0.16 [0.11, 0.22]** | **0.08 [0.00, 0.18]** | **0.36 [0.26, 0.48]** | **0.24 [0.13, 0.35]** |
| sd(Hemisphere) | **0.92 [0.70, 1.21]** | **0.71 [0.54, 0.95]** | **0.81 [0.62, 1.08]** | **0.98 [0.74, 1.32]** |
| cor(Intercept,time) | -0.07 [-0.44, 0.32] | -0.06 [-0.78, 0.74] | -0.18 [-0.52, 0.20] | -0.18 [-0.56, 0.26] |
| cor(Intercept,Hemisphere) | **-0.50 [-0.74, -0.19]** | **-0.66 [-0.84, -0.39]** | **-0.40 [-0.67, -0.05]** | -0.14 [-0.49, 0.22] |
| cor(Time,Hemisphere) | -0.24 [-0.59, 0.14] | -0.07 [-0.80, 0.75] | 0.19 [-0.18, 0.53] | -0.06 [-0.49, 0.36] |
| **Population-Level Effects** | | | | |
| Intercept | 0.34 [-0.24, 0.93] | **0.94 [0.52, 1.36]** | -0.02 [-0.61, 0.56] | -0.39 [-0.86, 0.08] |
| Time | 0.03 [-0.07, 0.12] | -0.08 [-0.19, 0.04] | 0.07 [-0.13, 0.27] | 0.01 [-0.15, 0.17] |
| Group(MG) | -0.15 [-0.97, 0.65] | -0.31 [-0.90, 0.28] | 0.04 [-0.78, 0.86] | 0.42 [-0.23, 1.06] |
| Hemisphere(Right) | **-0.60 [-1.07, -0.13]** | **-1.64 [-2.02, -1.25]** | 0.17 [-0.26, 0.59] | **0.48 [-0.03, 1.00]** |
| Time:Group(MG) | -0.07 [-0.21, 0.06] | -0.04 [-0.20, 0.13] | -0.17 [-0.44, 0.10] | -0.14 [-0.37, 0.09] |
| Time:Hemisphere(Right) | -0.02 [-0.09, 0.05] | -0.03 [-0.18, 0.12] | -0.01 [-0.11, 0.08] | -0.10 [-0.24, 0.05] |
| Group(MG):Hemisphere(Right) | 0.17 [-0.49, 0.84] | 0.28 [-0.26, 0.82] | -0.26 [-0.85, 0.33] | -0.11 [-0.83, 0.62] |
| Time:Group(MG):Hemisphere(Right) | 0.01 [-0.08, 0.12] | **0.17 [-0.04, 0.38]** | 0.01 [-0.12, 0.15] | 0.14 [-0.07, 0.35] |
| **Family-Specific Parameters** | | | | |
| sigma | 0.07 [0.05, 0.09] | 0.14 [0.11, 0.18] | 0.09 [0.07, 0.12] | 0.14 [0.11, 0.19] |
| **Posterior predictive check** | 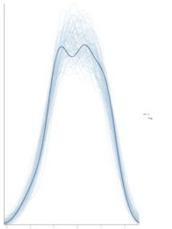 | 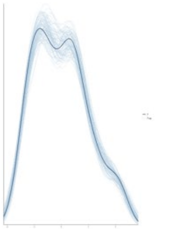 | 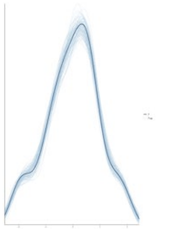 | 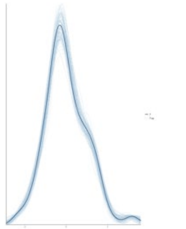 |

*Note.* The table shows group level, and population level effects, the z-score transformed values of the median estimate is followed by the credible intervals between brackets. The columns present each of the ROIs, namely, STG: Superior Temporal Gyrus; TTG: Transverse Temporal Gyrus; PCG: Precentral Gyrus; and PCL: Paracentral Lobule. SD indicates standard deviation, Cor indicates correlation between parameters with varying slopes. MG: Music Group. Effects that are pertinent to the hypotheses are in black, whereas all other effects are faded in grey. Effects that are credibly different from zero are in bold setting.

**Supplementary Materials S2:**

For the models that showed a three-way time-group-hemisphere interaction, we further probed the difference over time (pre and post training) within and between groups in each hemisphere. Table S4 shows the hypotheses tested for TTG surface area, including the mean estimate, credible intervals, and posterior probabilities for each test. We considered differences meaningful if credible intervals did not overlap 0 and had a posterior probability > 95%. Table S5 shows the same tests conducted for the model on STG cortical thickness.

Table S4. Group, and hemisphere differences interacting with time for TTG surface area.

| Outcome | Estimate | Est. Error | CrI. Lower | CrI. Upper | Evid. Ratio | Post. Prob |
|---|---|---|---|---|---|---|
| Time × Group × Hemisphere | 0.17 | 0.11 | 0.00 | 0.35 | 19.41 | 0.95 * |
| Time effect larger in RH than LH (Music) | 0.15 | 0.08 | 0.02 | 0.27 | 33.48 | 0.97 * |
| Time effect smaller in RH than LH (Control) | -0.03 | 0.07 | -0.15 | 0.09 | 1.84 | 0.65 |
| Music < Control overtime in LH | -0.04 | 0.08 | -0.17 | 0.10 | 2.04 | 0.67 |
| Music > Control over time in RH | 0.14 | 0.08 | 0.01 | 0.27 | 21.99 | 0.96 * |
| Decrease over time in Music–LH | -0.12 | 0.06 | -0.21 | -0.02 | 38.12 | 0.97 * |
| Decrease over time in Control–LH | -0.08 | 0.06 | -0.18 | 0.02 | 10.98 | 0.92 |
| Increase overtime in Music-RH | 0.03 | 0.06 | -0.07 | 0.13 | 2.30 | 0.70 |
| Decrease over time in Control–RH | -0.11 | 0.06 | -0.2 | -0.01 | 33.19 | 0.97 * |

Figure S1. Change in raw values of TTG's surface area, by group, hemisphere, and time point.

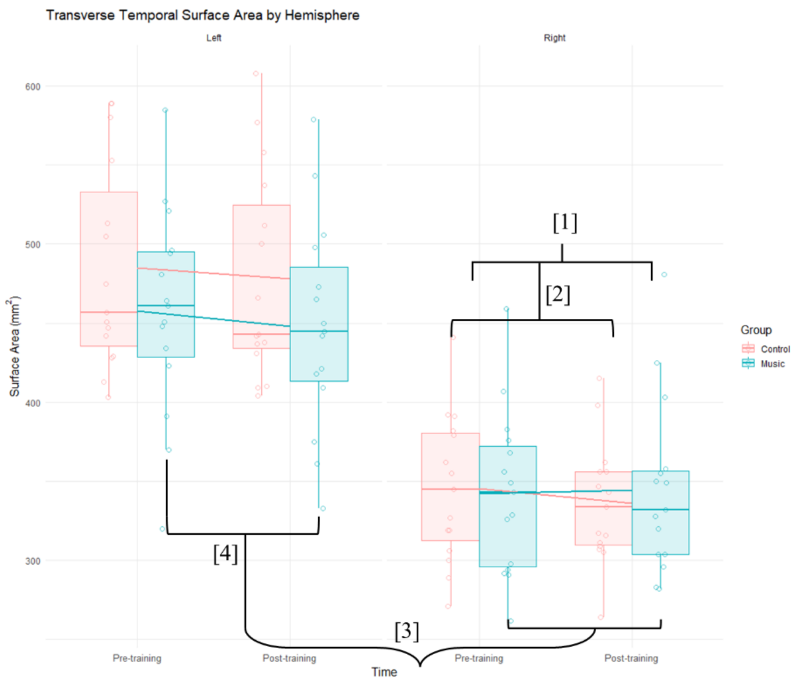


*Note.* Figure S1 shows the non-transformed raw values of TTG's surface area in $mm^2$. The hemispheres are shown in the left and right panels, respectively. The values for pre and post training are shown per group, the boxplot shows group averages and standard deviations in the error bars, where the data points are shown in the same color as the groups. The tests show that the three-way (time, hemisphere, group) interaction is evident in [1] the time effect between groups in the right hemisphere is larger in Music > Control [2] In the Control group, the right hemisphere decreases over time, [3] whereas in the Music group, the time effect is larger in the right > left hemisphere, and [4] the left hemisphere decreases over time.

Table S5. Group, and hemisphere differences interacting with time for STG cortical thickness.

| Outcome | Estimate | Est. Error | CrI. Lower | CrI. Upper | Evid. Ratio | Post. Prob |
|---|---|---|---|---|---|---|
| Time × Group × Hemisphere | -0.28 | 0.15 | -0.53 | -0.03 | 28.30 | 0.97 * |
| Time effect smaller in RH than LH (Music) | -0.17 | 0.11 | -0.34 | 0.01 | 15.05 | 0.94 |
| Time effect larger in RH than LH (Control) | 0.11 | 0.11 | -0.06 | 0.29 | 6.31 | 0.86 |
| Music > Control overtime in LH | 0.19 | 0.18 | -0.11 | 0.49 | 5.77 | 0.85 |
| Control > Music over time in RH | -0.09 | 0.18 | -0.40 | 0.21 | 2.35 | 0.70 |
| Increase over time in Music–LH | 0.11 | 0.13 | -0.10 | 0.32 | 4.01 | 0.80 |
| Decrease over time in Control–LH | -0.08 | 0.13 | -0.30 | 0.13 | 2.76 | 0.73 |
| Decrease overtime in Music-RH | -0.06 | 0.13 | -0.27 | 0.16 | 2.16 | 0.68 |
| Increase over time in Control–RH | 0.03 | 0.13 | -0.18 | 0.24 | 1.56 | 0.61 |

Figure S2. Change in raw values of STG's cortical Thickness, by group, hemisphere, and time.

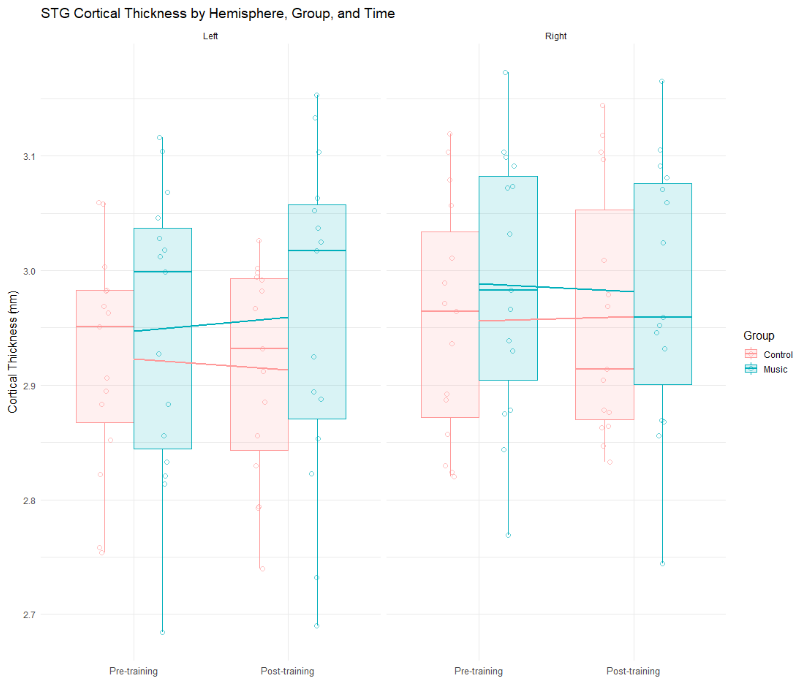


*Note.* Figure S2 shows the non-transformed raw values of STG's cortical thickness area in mm. The hemispheres are shown in the left and right panels, respectively. The values for pre and post training are shown per group, the boxplot shows group averages and standard deviations in the error bars, where the data points are shown in the same color as the groups. The hypotheses tests show that the three-way (time, hemisphere, group) interaction is evident, but no effects of time between groups in either hemisphere, or within-groups either within or between hemispheres.

### Supplementary Materials S3:

#### ***Brain-Behavior Relationships: Change in Motor Learning and Cortical Structures***

To assess brain-behavioral correlations, no outliers were identified in grey matter or behavioral indices, however, Shapiro-Wilk's test also revealed a violation of normality ($p < .05$). This was not improved with transformations, but visual inspection revealed acceptable normality, and as the behavioral outcomes are used in correlations, outliers do not directly affect the associations rather than within-variable distribution. For the behavioral outcomes, the homogeneity of variance is > .05 level for the change in sequence accuracy of the untrained sequence. Thus, Spearman (rather than Pearson) correlations were used, to take inhomogeneity of variances into account.

When examining associations between sequence accuracy and cortical measures of ROIs across all participants without Bonferroni correction, a negative association was found between sequence accuracy of untrained sequences and the surface area of the left PCG ($\rho = -0.43$, $p = .017$), which, when split up by group, was only found in the Music ($\rho = -0.582$, $p = .023$) and not in the Control group ($\rho = -0.136$, $p = .630$). Moreover, there was an inverse association between trained sequences' accuracy and surface area of the left PCG ($\rho = -0.53$, $p = .042$). Additionally, increase in sequence accuracy of trained sequences was inversely associated with surface area of both the PCL ($\rho = -0.540$, $p = .037$), and right PCL ($\rho = -0.522$, $p = .046$), see Table S6. These associations, although in unexpected direction, were only found in the Music and not in the Control group, however, none of these associations survived multiple comparison correction (*Bonferroni corrected* $\boldsymbol{\alpha} = 0.004$).

**Table S6.** Uncorrected significant correlations in the Music Group between changes in behavioral sequence accuracy and changes in surface area in grey matter ROIs

| Sequence | ROI | *ρ* | *p* | Corrected-*p* |
|---|---|---|---|---|
| Untrained | LH – PCG | -0.582 | 0.023* | 0.276 |
| Trained | LH – PCG | -0.531 | 0.042* | 0.504 |
| Trained | LH – PCL | -0.540 | 0.037* | 0.444 |
| Trained | RH – PCL | -0.522 | 0.046* | 0.552 |

*Note.* Change indices are derived from post-training minus pre-training values of sequence accuracy (the number of correctly performed sequences in 30 seconds) and surface area in ROIs indicated by LH: left hemisphere; RH: right hemisphere; PCG: precentral gyrus; PCL: paracentral Lobule. The tested sequences were Trained: sequences on which participants trained for four weeks; or Untrained: novel sequences only presented in time-points measurements. Spearman's *ρ* indicates the size of the correlation, significance at the $p < .05$ level is indicated by (*). None of these associations were significant in the Control Group.

**Figure S3.** Associations between change in motor performance and cortical structures.

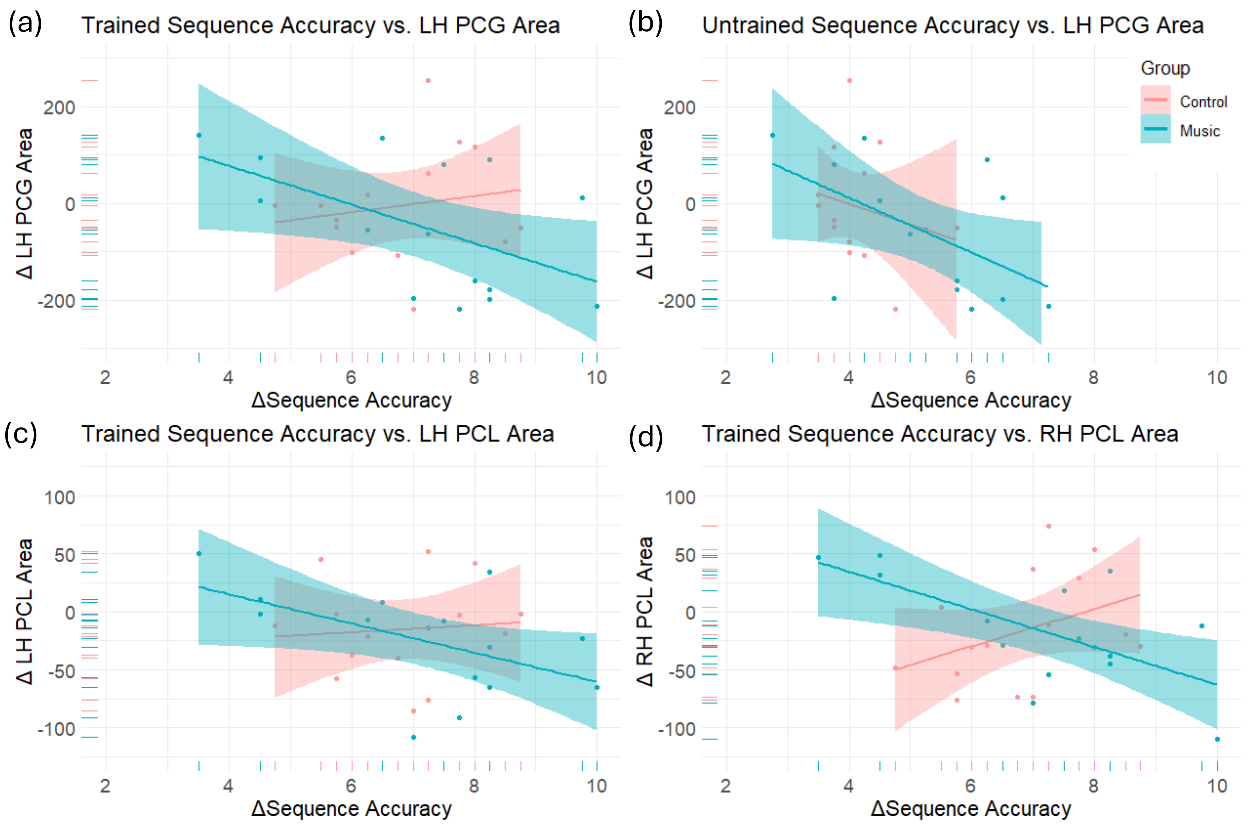


*Note:* Associations are displayed for the change in sequence accuracy and cortical structures between groups. Before Bonferroni correction, all correlations were significant for the Music but not the Control group. These correlations showed medium-sized effects but were no longer significant after multiple comparison correction.

Notably, all uncorrected significant associations are negative, and emerge only for motor areas, and only for the Music group. For PCL this is bilateral and only for trained sequences, and for PCG it is only in LH but for both sequences, suggesting this change to be related to general motor skill rather than sequence learning (albeit ipsilaterally to the trained hand). These findings, which as noted do not survive multiple correction comparisons but have medium-sized effects, suggest that PCG and PCL do change specifically based on musical cueing, but warrant further investigation in larger samples.

There are previous findings that support both ipsilateral and bilateral effects of motor leaning; for instance, finger movement tasks were previously shown to elicit ipsilateral functional activation of motor regions, including the motor cortex (Paek et al., 2014), while the left hemisphere is more specialized for motor sequences (Wiestler & Diedrichsen, 2013), and our behavioral findings indicate that sequence learning persisted after more general motor learning, measured through untrained sequences, had reached a ceiling. Moreover, while bilateral hemispheric involvement is observed even with single hand training, longer durations of training, leading to specialization, may diminish activity in the contralateral hemisphere over time, while ipsilateral effects remain (Wiestler & Diedrichsen, 2013).

### Supplementary Materials S4:

#### *Grey And White Matter Relationships*

In the Music group, a strong inverse relationship was found between FA of the left arcuate and volume of the left PCG (ρ = -0.604, *p* = .025), which was not significant after Bonferroni correction. The Control group showed strong positive associations in the right hemisphere between the arcuate fasciculus and volume of the PCL (*ρ* = .736, *p* = .006) and surface area of the PCG (*ρ* = .610, *p* = .030), which were also not significant after Bonferroni correction (*Bonferroni corrected **α*** = 0.004). Associations that were significant in the Music group were not significant for the Control group and vice versa, see Figure S4.

**Figure S4.** Associations between white matter connectivity and grey matter cortical structures.

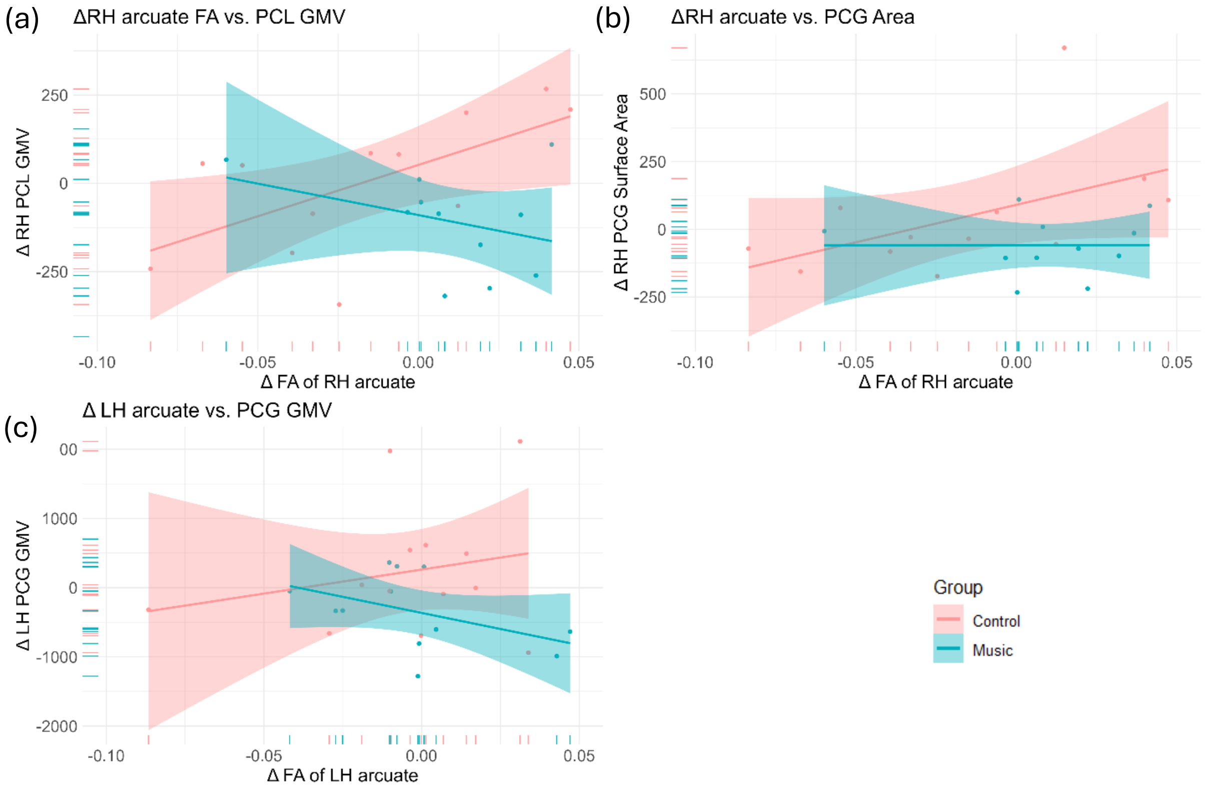


*Note:* Associations are shown between the change in the FA of the left arcuate fasciculus in relation the PCG volume for the Music group, and between the right arcuate fasciculus in relation to volume of the right PCL and area of the right PCG in the Control group. FA: fractional anisotropy; AF: arcuate fasciculus; Difference between post-training and pre-training is indicated by (Δ); LH: left hemisphere; RH: right hemisphere; PCG: precentral gyrus; PCL: paracentral lobule; SA: surface area; GMV: grey matter volume.

Here, the uncorrected findings indicate that for the Music group, the FA in the LH (ipsilateral to the trained hand) is negatively related to LH PCG volume, while for the Control group, RH FA (contralateral to the trained hand) is positively related to PCG surface area and PCL volume. As none of these results survive multiple comparison corrections, and interactions between FA values and grey matter indices are likely mostly physiological rather than based on the type of motor training, this is not interpreted further.
